# Conformational coupling of the sialic acid TRAP transporter HiSiaQM with its substrate binding protein HiSiaP

**DOI:** 10.1101/2023.03.04.531103

**Authors:** Martin F. Peter, Jan A. Ruland, Yeojin Kim, Philipp Hendricks, Jan Peter Siebrasse, Gavin H. Thomas, Ulrich Kubitscheck, Gregor Hagelueken

**Affiliations:** Institute of Structural Biology, University of Bonn, Venusberg-Campus 1, 53127 Bonn, Germany; Clausius Institute for Physical und Theoretical Chemistry, University of Bonn, Wegelerstr. 12, 53127 Bonn, Germany; Department of Biology (Area 10), University of York, York YO10 5YW, United Kingdom

## Abstract

The tripartite ATP-independent periplasmic (TRAP) transporters use an extra cytoplasmic substrate binding protein (SBP) to transport a wide variety of substrates in bacteria and archaea. The SBP can adopt an ‘open’ or ‘closed’ state depending on the presence of substrate. The two transmembrane domains of TRAP transporters form a monomeric elevator whose function is strictly dependent on the presence of a sodium ion gradient. Insights from experimental structures, structural predictions and molecular modeling have suggested a conformational coupling between the membrane elevator and the substrate binding protein. Here, we use a disulfide engineering approach to lock the TRAP transporter HiSiaPQM from *Haemophilus influenzae* in different conformational states. The SBP, HiSiaP, was locked in its substrate-bound form and the transmembrane elevator, HiSiaQM, was locked in either its predicted inward- or outward-facing states. We characterized the disulfide-locked variants and used single-molecule total internal reflection fluorescence (TIRF) microscopy to study their interactions. Our experiments demonstrate that the SBP and the transmembrane elevator are indeed ‘conformationally coupled’, meaning that the open and closed state of the SBP recognize specific conformational states of the transporter and vice versa.

## Introduction

Tripartite ATP-independent periplasmic (TRAP) transporters are widely distributed in bacteria and archaea, and are also found in common pathogens ^1,2^. Together with the ATP-binding cassette (ABC) importers and the tripartite tricarboxylate transporters (TTT), they define the three classes of substrate binding protein (SBP)-dependent transporters. The SBP (commonly referred to as the P-domain in TRAP transporters) is a soluble protein that freely diffuses in the periplasm of Gram-negative bacteria or is associated with the cell membrane in Gram-positive bacteria and archaea ^3^. It scavenges substrate molecules and delivers them to the membrane transporter. It is believed that the SBP is advantageous in situations where low substrate concentrations are encountered and that it serves as a substrate store ^4^.

The first structural information about TRAP transporters was provided by two high-resolution crystal structures of the P-domain of the sialic acid (Neu5Ac) TRAP transporter HiSiaPQM from *Haemophilus influenzae* in both the apo and holo states ^5,6^. Since then, many more structures have been determined, confirming the general architecture of the P-domain as two globular lobes connected by a long backbone helix (Fig. 1, red). Substrate binding induces a bend in this helix, closing the two lobes around the substrate molecule. This motion is often compared to that of a Venus flytrap and has been studied in detail, not only with high-resolution structures, but also with biophysical techniques and molecular dynamics (MD) simulations ^7–12^. In contrast, structural information on the two transmembrane domains, i.e., the smaller Q-domain with 4 transmembrane (TM) helices and the larger M-domain with 11 TM helices (Fig. 1 blue, yellow), has only recently been obtained. Two cryo-EM structures revealed that TRAP transporters are elevator-type transporters (reviewed in ^13^), but with a striking twist: TRAP transporters are monomeric elevators ^14,15^. The key to this unusual feature is the Q-domain, the function of which has been a mystery for a long time. The structures revealed that its four TM helices form a unique helical sheet that wraps around the M-domain and serves to enlarge the stator portion of the latter. This is thought to anchor the stator domain in the membrane, allowing the elevator portion to move up and down, yielding the outward- and inward-facing states of the transporter (OFS and IFS, respectively). How the conformationally flexible SBP interacts with these different states of the moving elevator transporter is an interesting structural question. Initial insights have come from a combination of experimental structures, structural predictions using the AlphaFold2 algorithm and repeat swap modeling ^14,16,17^. It was predicted that the closed-state SBP forms a complex with the IFS of the elevator and that the open-state SBP structurally matches and interacts with its OFS. While the predicted binding interfaces could be confirmed by mutagenesis ^14^, a direct observation of this conformational coupling would support the prediction and significantly improve our understanding of TRAP transporters.

**Fig. 1:**
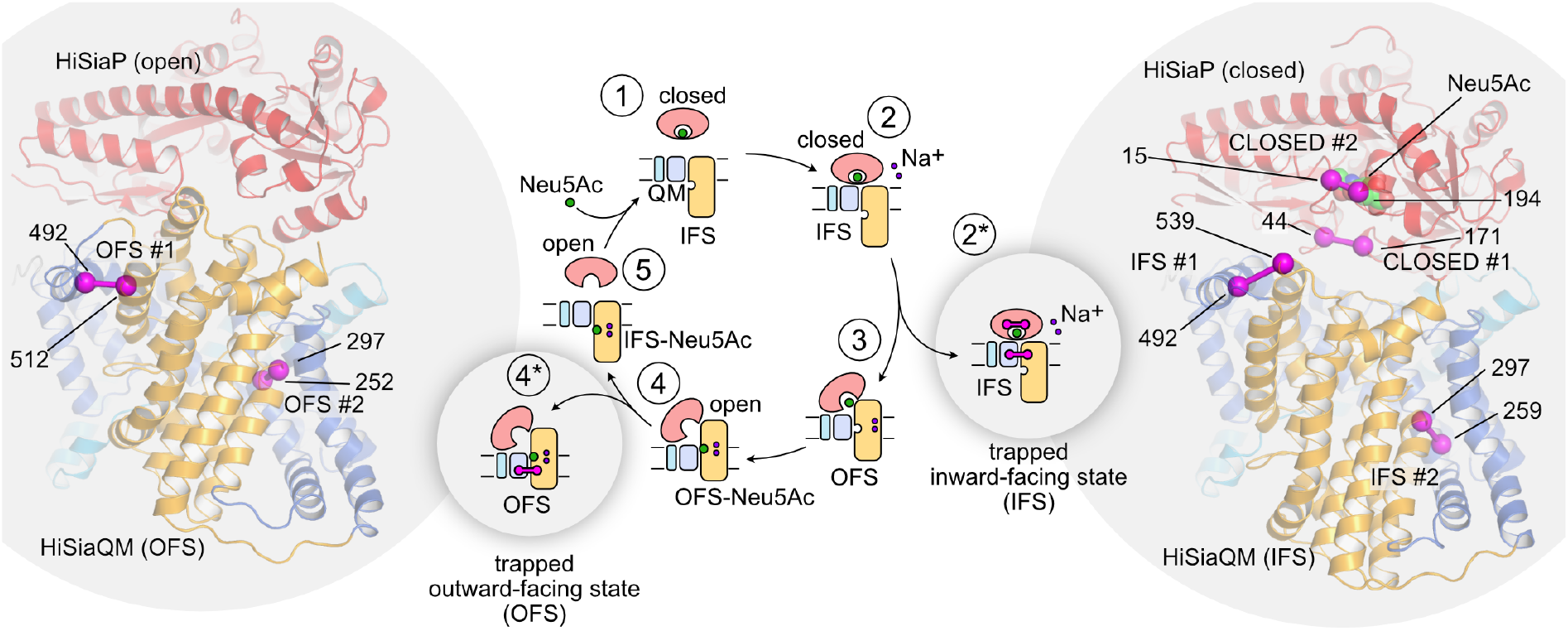
Trapping a TRAP transporter. The transport cycle of TRAP transporters as described in ^13^ is depicted schematically in the middle of the figure. To trap the transporter in specific states, the stator and elevator domains were cross-linked by disulfide engineering to yield states 2^*^ and 4^*^. The cartoon representations of HiSiaPQM on the left and right of the figure are AlphaFold2 models of the tripartite complex ^14^. The individual domains of the transporter are colored as in the schematic (HiSiaP: red, HiSiaQM stator: blue, HiSiaQM elevator: yellow). Residues that were mutated to cysteines to form disulfide bonds are indicated by magenta spheres and the disulfide bond is shown as a magenta bar connecting the two residues. In state 2^*^, the bound sialic acid is shown as spheres.

Here, we used disulfide engineering techniques ^18,19^ to trap the three domains of the sialic acid TRAP transporter HiSiaPQM in its major conformational states, i.e., a permanently closed state of the P-domain, as well as the IFS and OFS of the elevator domains. We used X-ray crystallography and biophysical approaches to characterize the conformationally trapped domains. We then combined the different trapped domains and used single molecule total internal reflection fluorescence (TIRF) microscopy to study their interactions.

## Results

### Design of a conformationally trapped TRAP transporter

We searched the structural model of the HiSiaPQM complex for sites where the different domains can be locked in place by intra-chain disulfide bonds. As shown in Fig. 1 (right), we selected two pairs of residues in HiSiaP (S44C/S171C and S15C/A194C) and named the two variants HiSiaP CLOSED #1 and HiSiaP CLOSED #2, respectively. For both variants, the thiol groups of the introduced cysteines would likely be in close proximity in the substrate-bound state (Supplementary Fig. 1), and the formation of a disulfide bond would therefore be expected to lock the SBP in this conformation. We have previously shown with various techniques that HiSiaP does not visit the closed conformation in the absence of sialic acid ^9,10,20^, allowing us to simply use the apo protein in experiments addressing the open state of HiSiaP. For HiSiaQM, we selected four pairs of residues to be converted to disulfides. The pairs A492C/Q539C and M259C/M297C should lock the elevator domain in its IFS (HiSiaQM IFS #1 and HiSiaQM IFS #2, respectively) (Fig. 1, right, Supplementary Fig. 1), while the pairs A492C/L512C and Y252C/M297C would trap the OFS (HiSiaQM OFS #1 and HiSiaQM OFS #2, respectively) (Fig. 1, left, Supplementary Fig. 1). The selection was guided by our previously published structural models ^14^ and inspired by the residues selected by Mulligan et al. ^19^ to trap the structurally related VcINDY elevator transporter in its IFS and OFS.

### Disulfide linked HiSiaP mutants bind Neu5Ac and can be trapped in the closed state

The CLOSED #1 and #2 variants of HiSiaP (Fig. 1) were expressed in *E. coli* and could be purified analogously to the wild-type protein. To confirm that the double cysteine mutants were still able to bind Neu5Ac, we used isothermal titration calorimetry (ITC) and determined a dissociation constant (K_D_) of 61 nM for the wild-type HiSiaP protein (Fig. 2), in agreement with previous studies ^12,14^. HiSiaP CLOSED #1 clearly bound sialic acid, but with an approximately 13-fold weaker affinity (K_D_ = 0.81 μM). It is known that even very conservative changes of the residues in the Neu5Ac binding pocket can lead to such changes in the K_D_^7^. The second variant, HiSiaP CLOSED #2, showed a different behavior in the ITC experiments (Fig. 2c). The titration curves of this variant consistently had a biphasic shape and could be fitted with two independent binding reactions, one with picomolar affinity (K_D_ of ∼2 pM) and one with micromolar affinity (K_D_ of ∼2 μM, Fig. 2c). We hypothesized that this observation was due to a fraction of HiSiaP CLOSED #2 forming a disulfide bond upon binding Neu5Ac, resulting in a very slow k_off_ rate and hence a small K_D_. This experiment also suggested that the presence of the cysteines alone, in the absence of an oxidizing agent, results in only a small fraction of the HiSiaP CLOSED state. To investigate this hypothesis, we used nanoDSF to follow the ligand binding process while varying the presence of ligand and oxidant (schematic of experimental design shown in Fig. 2d).

**Fig. 2:**
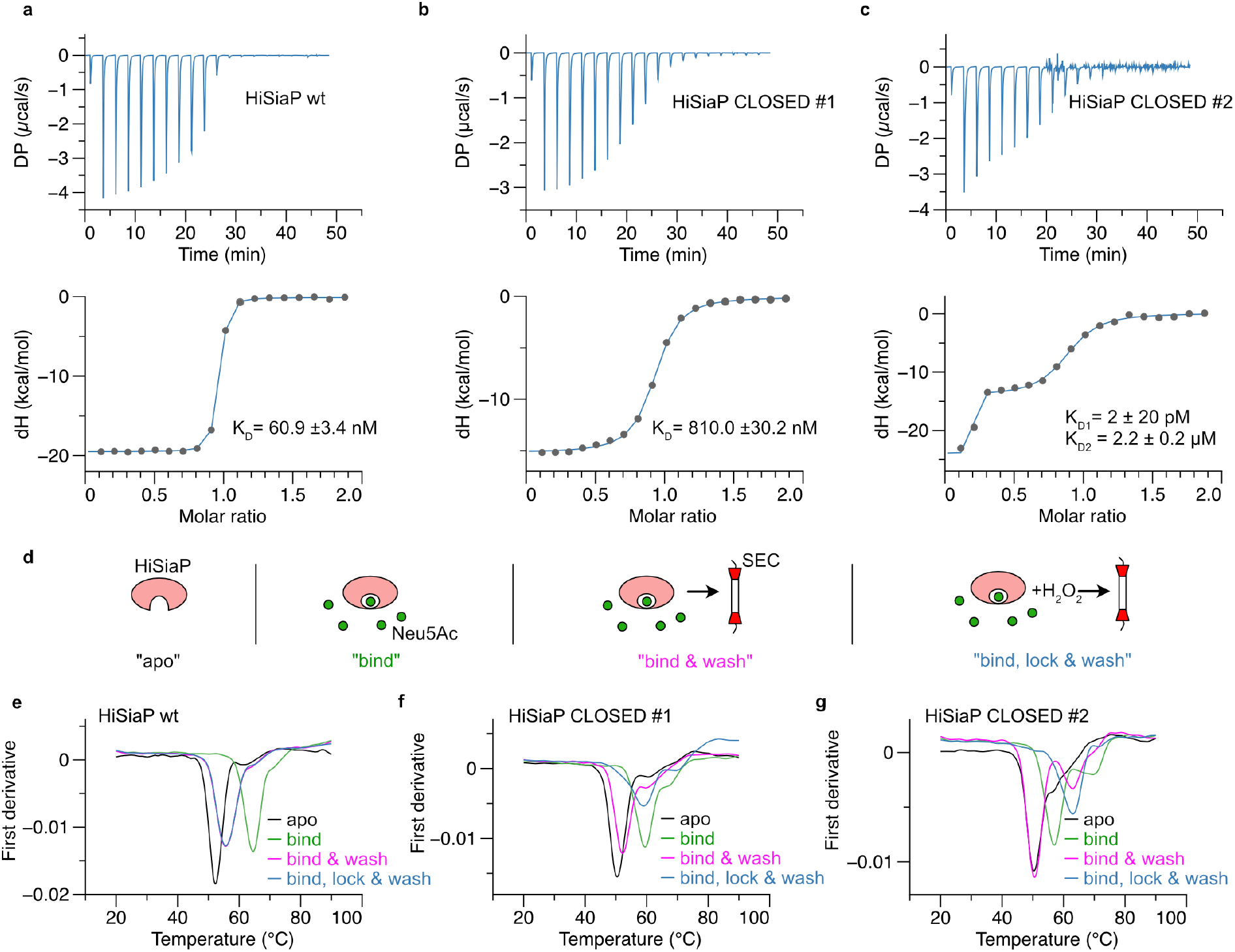
Trapping HiSiaP in its Neu5Ac-bound closed state. **a-c)** Binding of Neu5Ac to HiSiaP wt and its CLOSED #1 and #2 variants followed by ITC. The titration curves are shown on the top, the binding isotherm at the bottom of each panel. **d)** Schematic illustrating the sample preparation for the nanoDSF experiments in panels e-g. **e-g)** nanoDSF experiments with apo HiSiaP (black), HiSiaP in the presence of 1 mM Neu5Ac (green), the HiSiaP/Neu5Ac complex following ligand removal by gel filtration (magenta) and the H_2_O_2_- and then gel filtration-treated sample (blue). The curves represent the first derivative of the raw melting curves.

As expected, binding of sialic acid to wild-type HiSiaP resulted in a significant thermal stabilization of the protein with a ∼12°C shift in its melting temperature from ∼50°C to ∼62°C (Fig. 2e, black and green curves). For the HiSiaP CLOSED #1 and #2 variants, the addition of sialic acid also clearly stabilized the proteins, but not as much as observed for the wild type (black vs. green curves in Fig. 2e-g). In the next set of experiments, we incubated the proteins with sialic acid and then removed the excess sialic acid by gel filtration (Fig. 2d “bind & wash” and magenta curves in Fig. 2e-g). While the wild-type protein showed a slightly elevated melting temperature (Fig. 2e, magenta curve (covered by blue curve)), we found biphasic melting curves for the HiSiaP CLOSED #1 and #2 variants, where the largest fraction had melting temperatures very similar to the samples without any Neu5Ac added. The smaller fraction, however, had an increased melting temperature (compare the magenta and green curves in Fig. 2f, g), which was particularly evident for HiSiaP CLOSED #2. This was consistent with the observation made in the ITC experiments (Fig. 2c) and supports the notion that a portion of HiSiaP CLOSED #2 spontaneously formed a disulfide bond. To increase this fraction, we added H_2_O_2_ to the Neu5Ac binding step of the experiment and removed the oxidant together with the excess Neu5Ac by gel filtration (Fig. 2d, “bind, lock & wash”). Strikingly, the nanoDSF analysis now showed a single inflection point with a melting point significantly higher than that of the apo form of the two variants, demonstrating that the SBPs had been successfully locked in their closed state (Fig. 2e-g, blue curves).

### A crystal structure of HiSiaP trapped in its closed state

To verify that the disulfide-linked SBPs had the expected structure, we set up crystallization experiments of the H_2_O_2_-treated HiSiaP CLOSED **#**1/2 variants. Heavily intergrown crystals of HiSiaP CLOSED **#**2 were obtained and single crystalline pieces could be broken off for diffraction data collection at the PETRA III synchrotron (DESY, Hamburg, Germany). A 1.9 Å data set (space group I222) was collected and the structure was solved by molecular replacement using the closed state of HiSiaP as a search model (PDB-ID: 3B50 ^6^). A clear solution was found (Fig. 3a) and inspection of the electron density map revealed density at the expected position of the disulfide bond (Fig. 3b) and the sialic acid binding site (Fig. 3c), demonstrating that the disulfide bond had indeed been formed and the sialic acid molecule had been captured in its binding site. The structure was refined to R/R_free_ factors of 0.211/0.259 and good geometric parameters as determined by MolProbity ^21^ (Supplementary Table 1). Fig. 3a shows an overlay of the HiSiaP CLOSED #2 structure with that of the closed-state of the wild-type protein (PDB-ID: 3B50 ^6^). The two structures overlap with an r.m.s.d. value of 0.24 Å over 300 C_α_ atoms, and except for the introduced disulfide bond, there is no significant change between the backbone or side chain atoms of the two structures. Even the positions of the visible water molecules in the binding site are conserved (not shown). We found that traces of a second conformation of C194 were visible in the electron density (Fig. 3b). In the light of the above experiments, we conclude that this is most likely due to reduction of the disulfide bond due to radiation damage ^22^.

**Fig. 3:**
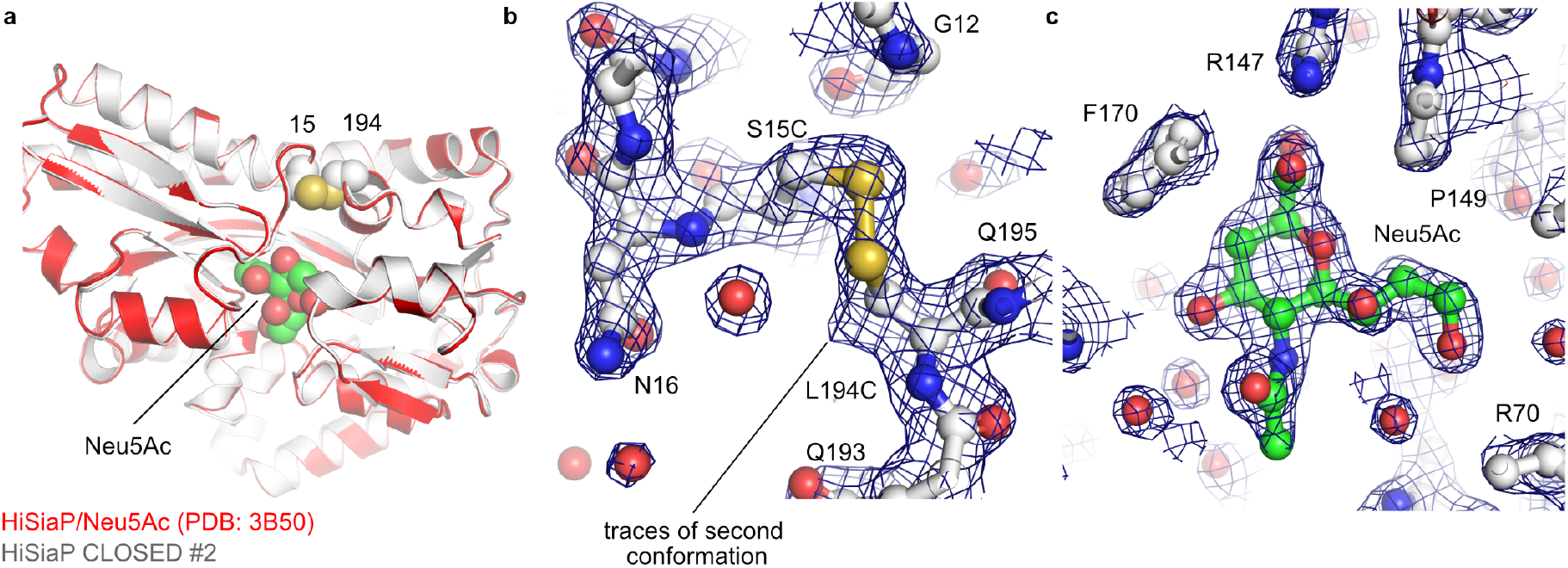
Crystal structure of HiSiaP 15C/194C (CLOSED #2). **A)** A superposition of the closed-state crystal structure of HiSiaP (red) in complex with Neu5Ac (PDB-ID: 3B50 ^6^) and the disulfide linked HiSiaP CLOSED #2 variant from this study (white). The two introduced cysteines at positions 15 and 194 are shown as spheres. **b)** Detail of the disulfide bond with 2F_o_-F_c_ density contoured at 1.5 σ. The map shows a fraction of the cysteine at position 194 populates a different rotamer and is hence not involved in a disulfide bridge. The positions of selected residues are indicated. **c)** 2F_o_–F_c_ density map (contoured at 1.5 σ) of the bound Neu5Ac molecule. The locations of selected residues are indicated.

### Disulfide-linking the elevator domain of HiSiaQM

We then expressed and purified the HiSiaQM IFS/OFS double cysteine variants shown in Fig. 1 and Supplementary Fig. 1. In each case, the protein was purified similarly to the wild type, while any disulfide linked oligomers were removed by gel filtration. To test experimentally, whether the designed disulfide bonds were actually present, we used single-molecule TIRF microscopy and the AF555-labeled VHH_QM_3 ^14^, to probe the conformational state of the disulfide-bonded QM-domains. As known from our cryo-EM structure (PDB-ID 7QE5), VHH_QM_3 binds on the periplasmic side of HiSiaQM, at the boundary between the stator domain and the inward-facing elevator domain with a very high affinity of 0.64 nM. Thus, if the disulfide bonds in the QM-domain formed as intended, VHH_QM_3 should still bind to the HiSiaQM IFS #1/2 variants but likely not to the OFS #1/2 variants. We incorporated all four mutants into DOPC-based solid supported bilayers (SSBs) and added 1 mM H_2_O_2_ during the reconstitution procedure to ensure that the disulfide bond was formed as soon as the engineered disulfides contacted each other. We ensured that the H_2_O_2_ concentration did not exceed 1 mM, since we observed aggregation of the QM-domain/nanobody complex in the bilayer at H_2_O_2_ concentrations higher than 2 mM.

As previously reported ^14^, VHH_QM_3 bound irreversibly to the immobilized wild-type HiSiaQM (Fig. 4a). A similar level of interaction was observed for HiSiaQM IFS #1 (Fig. 4b). The HiSiaQM IFS #2 mutant also interacted with VHH_QM_3, but the number of binding events observed was significantly lower, presumably indicating a slightly different conformation of HiSiaQM IFS #2 (Fig. 4b). For the HiSiaQM OFS #1/2 variants, only very spurious interactions with VHH_QM_3 were detected (Fig. 4c). Strikingly, the binding of the nanobody to HiSiaQM OFS#1/2 could be restored when H_2_O_2_ was omitted from the experiment. While these experiments strongly suggested that the disulfide bonds had been formed in the lipid bilayer, the experiment did not exclude the possibility that HiSiaQM OFS#1/2 were trapped in a conformation that did not represent a productive state of the transporter. However, subsequent experiments strongly suggested that the QM-domain retains its functional integrity (see below).

**Fig. 4:**
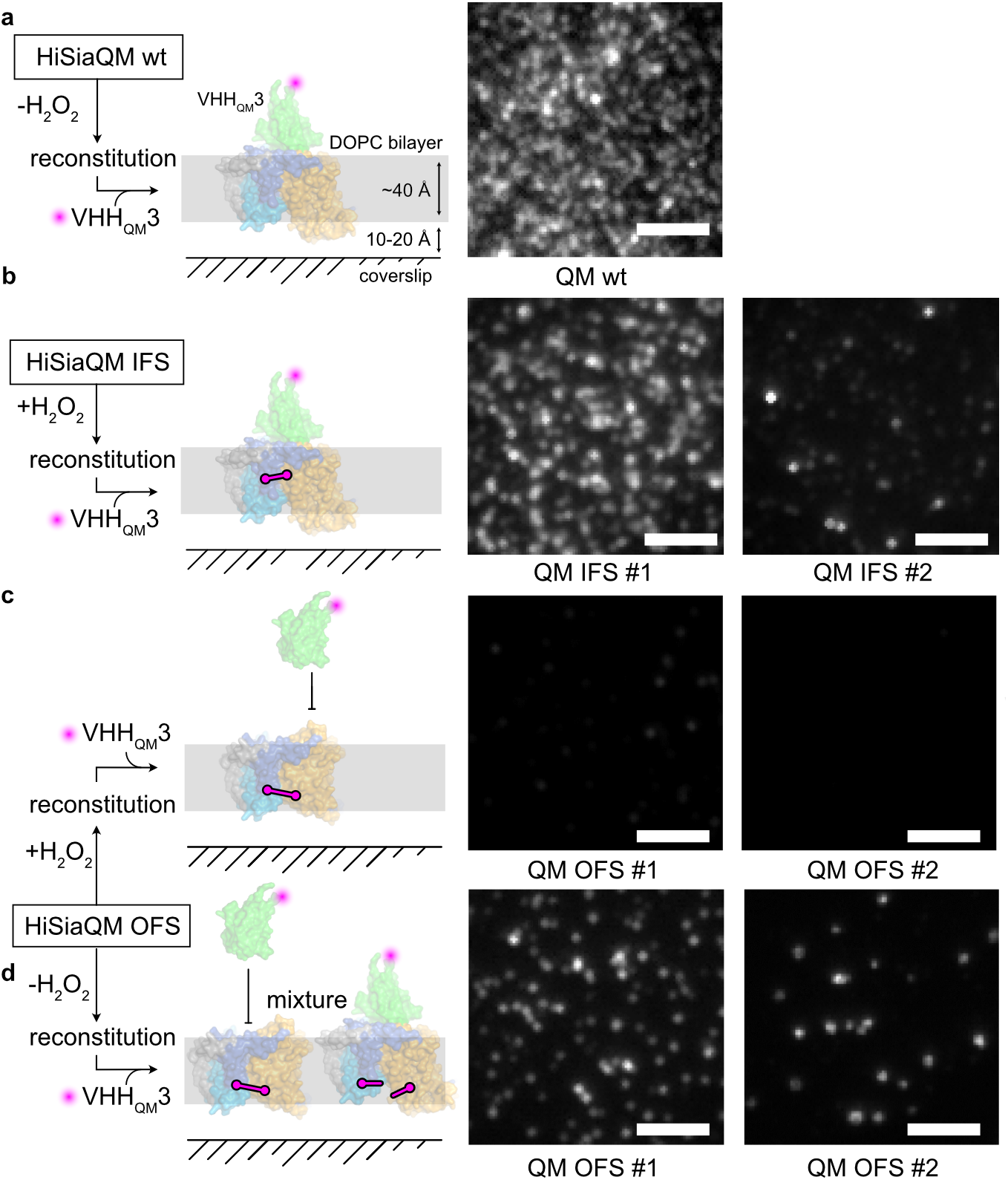
VHH_QM_3 binding to the IFS/OFS #1/2 variants of HiSiaQM. **a)** The schematic on the left depicts the setup of the single molecule TIRF experiment. The micrograph on the right shows that VHH_QM_3 interacts with the wild-type QM-domains in the bilayer. **b)** Same as a) but with HiSiaQM IFS #1/2. **c)** Same as a), but with HiSiaQM OFS #1/2. **d)** Same as c) but without adding H_2_O_2_ during the reconstitution. The scale bars equal 3 μm

Interestingly, we noted that for the DDM detergent solubilized transporter, the addition of 1 mM H_2_O_2_ was not sufficient to lock the HiSiaQM OFS#1/2 variants, as evident from gel filtration experiments, were VHH_QM_3 still bound to the H_2_O_2_-treated transporter (Supplementary Fig. 2). A possible explanation would be that the elevator domain is not sufficiently mobile in DDM and never or at least very rarely visits the upward facing conformation.

### Interactions of the disulfide-linked P- and QM-domains observed by TIRF microscopy

The single molecule TIRF microscopy setup was then used to investigate the effect of disulfide trapping on the interactions between HiSiaQM and HiSiaP. The first set of experiments was performed with wild-type HiSiaQM, HiSiaQM IFS #1, HiSiaQM OFS #1 and AF647-labeled HiSiaP in the presence of sialic acid. As expected ^14^, we observed many interactions between the wild-type P-domain and the wild-type QM-domains (Fig. 5a). A slightly lower level of interactions was observed between the wild-type P-domain and HiSiaQM IFS #1 (Fig. 5b). In contrast, the HiSiaQM OFS #1 mutant showed a significantly lower (−50 %) rate of interactions with HiSiaP under these conditions (Fig. 5c).

**Fig. 5:**
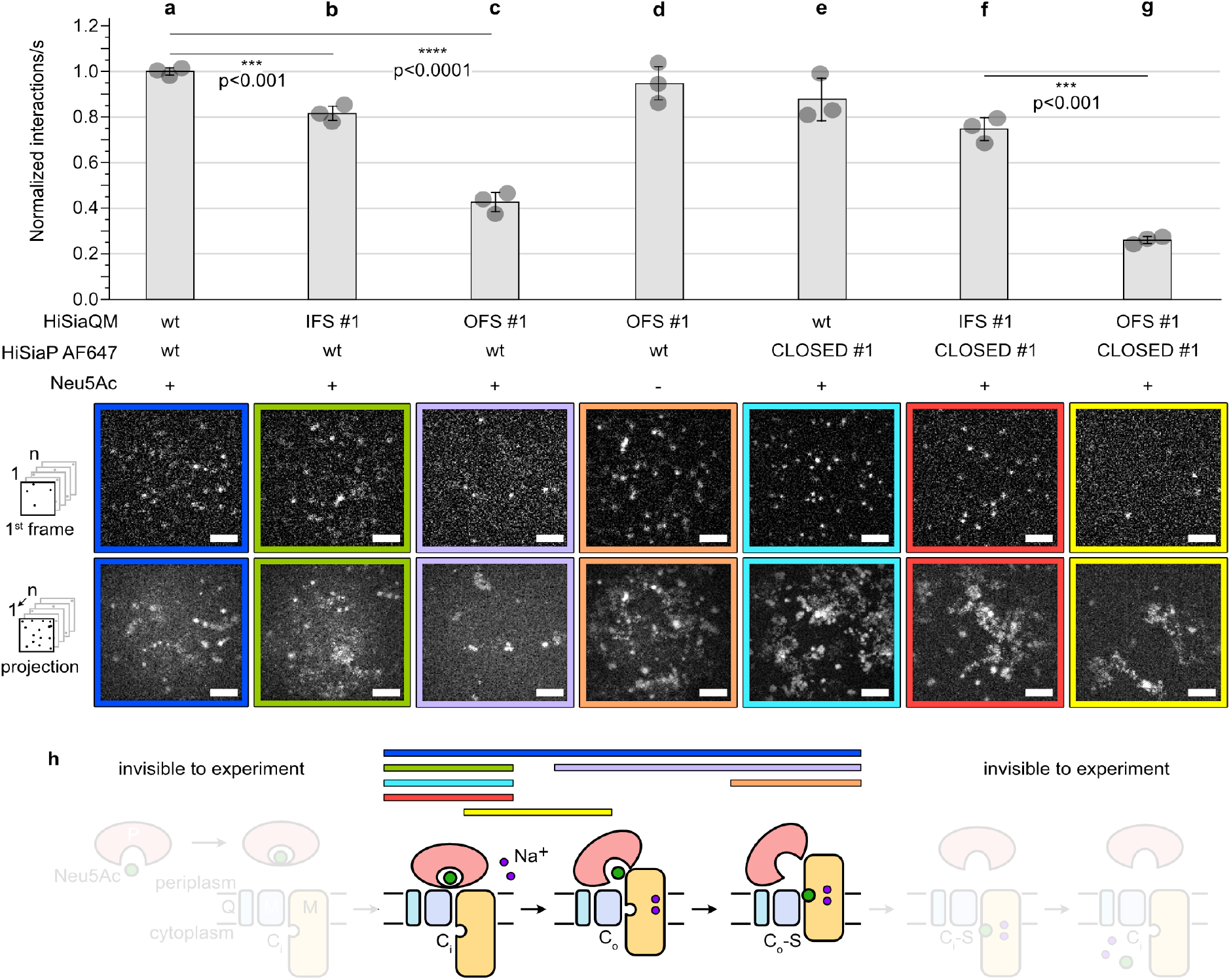
Single molecule TIRF microscopy of trapped TRAP transporter domains. **a-g)** Top: Normalized interactions per second between AF647 labelled HiSiaP variants and the indicated HiSiaQM constructs. Bottom: first frame of an image sequence of a typical set of data and the corresponding maximum intensity projection of the respective image sequence. A movie with all conditions can be found in the supplementary information (Supplementary Movie 1). **h)** In the schematic, colored horizontal bars mark the likely state of the transporter observed in the experiments a-g). The statistical significance of differences between selected experiments was assessed by applying a two-sided unpaired Student’s t-test with a 95% confidence interval. The scale bars equal 3 μm.

In the next experiment, HiSiaQM OFS #1 was combined with wild-type HiSiaP in the absence of sialic acid, conditions under which the latter does not visit its closed state ^9,10,20^. Strikingly, the open SBP interacted with HiSiaQM OFS #1 to a similar extent as observed for the two wild-type proteins in the presence of sialic acid (Fig. 5d), in line with the model shown in Fig. 1 (left). Further, the observed interaction between apo HiSiaP and HiSiaQM OFS #1 strongly indicates that the latter is structurally intact.

In the next step, HiSiaP CLOSED #1 was added to the mix. As expected from the wild-type experiment in Fig. 5a and our crystal structure (Fig. 3), interactions with wild-type HiSiaQM were observed (Fig. 5e). The interaction of HiSiaP CLOSED #1 with HiSiaQM-domains IFS #1/OFS #1 are shown in Fig. 5f, g. Clearly, HiSiaP CLOSED #1 preferably interacted with the IFS #1 variant of HiSiaQM. Taken together, these experiments confirm the expectation that the closed state of HiSiaP preferentially interacts with the IFS of HiSiaQM and the open state HiSiaP preferentially interacts with the OFS of HiSiaQM. The same set of experiments was conducted with HiSiaQM OFS#2/IFS #2 and HiSiaP CLOSED #2, leading to very similar results (Supplementary Fig. 3).

The SSB TIRF setup allowed us to add soluble components during an ongoing experiment and to observe the effects in real time. This led us to design an experiment that provided further evidence that the engineered disulfide bonds were indeed formed in the lipid bilayer. Fig. 6a shows such an experiment, where the HiSiaQM OFS #1 variant was incorporated into the SSB in the absence of sialic acid and in the presence of the maleimide-AF647 labeled P-domain. In this case, HiSiaQM OFS #1 was not pre-treated with H_2_O_2_ and therefore, only a fraction of the engineered disulfide bonds was likely formed. Hence, the number of interactions between the open state HiSiaP and HiSiaQM OFS #1 was lower than observed in the experiment shown in Fig. 5d. We then added H_2_O_2_ to induce disulfide bond formation and allowed the system to equilibrate for 10 min. Indeed, a significant increase in the observed interactions clearly indicated that the disulfide bonds had been formed (Fig. 6b). We then added the reducing agent DTT (dithiothreitol) and allowed the system to equilibrate again for 10 min. The number of interactions was now greatly reduced and lower than at the beginning of the experiment, presumably, because after the addition of DTT any preformed disulfide bonds were now broken and most of the HiSiaQM proteins were in the IFS (Fig. 6c).

**Fig. 6:**
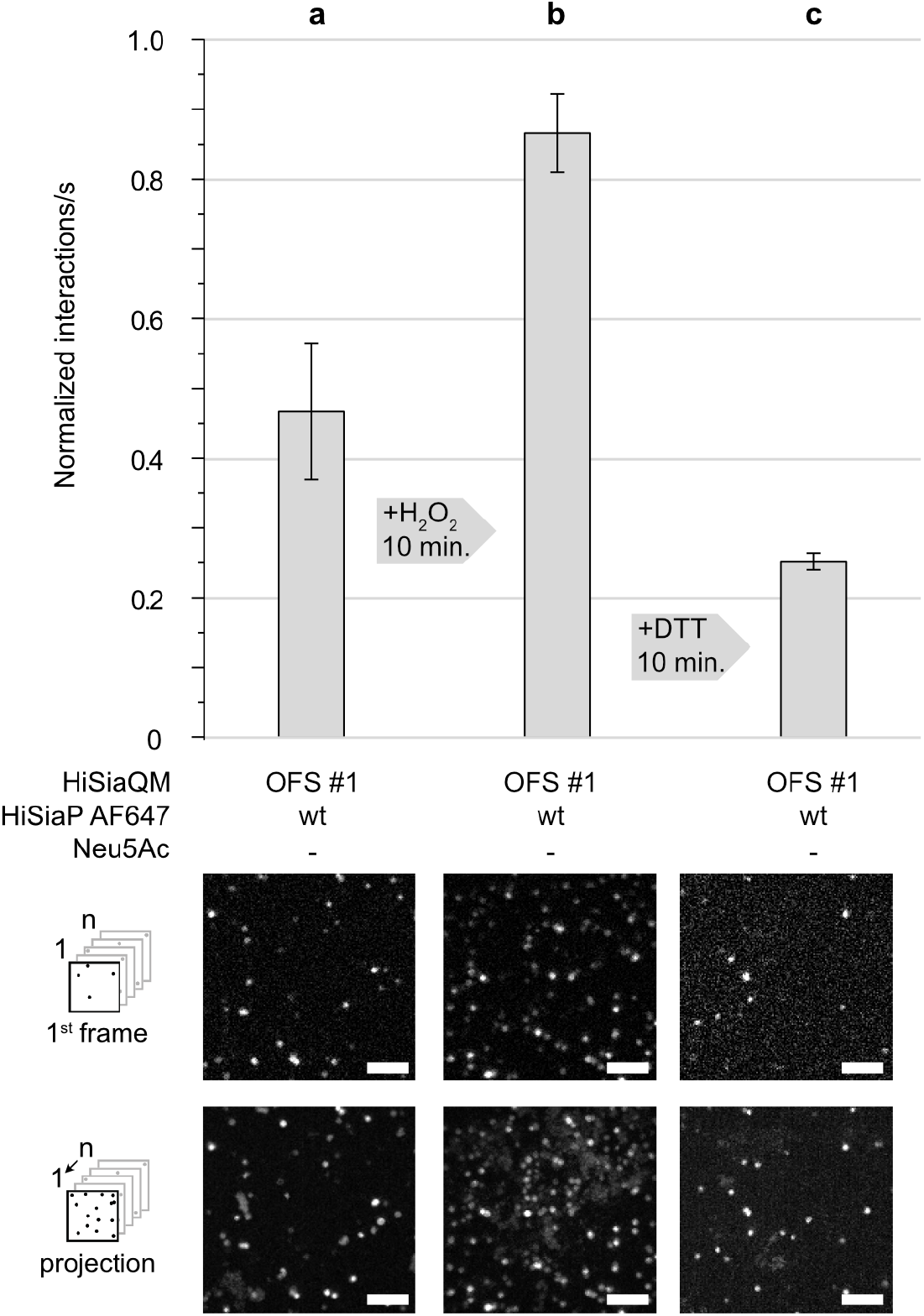
Following disulfide formation in HiSiaQM OFS #1 in real time. **a)** HiSiaQM OFS #1 was not pre-treated with H_2_O_2_ and incorporated into DOPC SSBs and interactions with the AF647 labelled P-domain were observed in the absence of sialic acid. **b)** H_2_O_2_ was added to induce disulfide bond formation, leading to an increased rate of interactions between the P- and QM-domains. **c)** This could be reversed by reducing the disulfide bonds with DTT (Supplementary Movie 2). The scale bars equal 3 μm.

During the SSB TIRF experiments, we observed a peculiar behavior of the AF647-labeled P-domains, especially of the HiSiaP CLOSED #1/2 mutants (Fig. 5e-g): A significant fraction of highly mobile SBPs appeared in the focal plane of the microscope and stayed in the membrane plane for a longer time than observed for the wild-type P-domain (Fig. 5a). This effect can be seen by comparing the projection images in Fig. 5a-g (bottom row of micrographs) with the single frame images (top row of micrographs). The fast motion leads to the formation of a cloudy background in the projection images. A possible explanation for this observation is that under the corresponding experimental conditions, the three domains of the transporter are in “compatible” conformational states, i.e., after dissociation, a locked closed P-domain will more quickly find a QM-domain that matches its conformational state, allowing a new interaction to form on the membrane and thus a longer microscopic observation. This was particularly evident for the HiSiaP CLOSED #1 and #2 variants, but was also visible for apo HiSiaP in combination with HiSiaQM OFS #1 (Fig. 5d, Supplementary Movie 1, 2).

## Discussion

The recent advances of the structural knowledge of TRAP transporters have enabled us to engineer “locked” P- and QM-domains by disulfide engineering ^18^. Our high-resolution crystal structure of HiSiaP CLOSED #2, together with ITC and nanoDSF data, demonstrated that our efforts had the intended effect on the P-domain (Fig. 2, 3). For the QM-domains, the TIRF microscopy experiments with VHH_QM_3 and apo HiSiaP provided evidence that also here, the engineered disulfide bonds were formed and that the QM-domains still had their native structure (Fig. 5, 6).

By combining locked variants and wild-type proteins in the presence or absence of Neu5Ac, we were able to perform single-molecule TIRF microscopy experiments on different states of the transport cycle (Fig. 5h). Our experiments strongly support a key postulation of the current TRAP transporter working hypothesis ^14^. That is, the closed state P-domain preferentially interacts with the inward facing elevator and the open state P-domain (i.e., the apo P-domain) preferentially interacts with the outward facing elevator.

Unexpectedly, the locked P-domain still interacted with the OFS of the transporter to a small degree (Fig. 5g). While a simple structural superimposition of the C-terminal lobe of the closed P-domain onto a model of HiSiaQM in its OFS leads to clashes ^14^, small conformational rearrangements could render the interaction possible, and an intermediate between states 2 and 3 of the transport cycle would be a candidate to explain the observed state (Fig. 1, Fig. 5f). Another explanation would be just one lobe of the P-domains interacting with one domain (stator or elevator) of the QM-domain, while the other binding interface is not formed. This would presumably result in a lower affinity explaining the smaller number of identified interactions.

Our observation that the engineered disulfide bonds of the HiSiaQM IFS/OFS constructs were readily formed in the DOPC bilayer, indicates that the transporter visits both the IFS/OFS of the transport cycle, even in the absence of both a Na^+^-gradient and the P-domain. A similar observation has been made for the GltPh transporter ^23,24^. While it is interesting that HiSiaQM appears to behave quite differently in detergent, it is well known that the structure and function of membrane proteins can be strongly influenced by the lipid environment ^25^. The role of the lipid environment for elevators was highlighted in a recent MD study on GltPh, which indicated that the switch between the IFS and OFS has a strong influence on the lipid bilayer ^26^.

In previous experiments in which the transporter was reconstituted in liposomes in the absence of both a Na^+^-gradient and the P-domain, no transport was observed ^27^. Thus, simple up and down movement of the transporter appears to be insufficient to drive transport. A possible explanation for this conundrum would be an influence of both the P-domain and the Na^+^-gradient on the affinity between the QM-domain and its substrate, Neu5Ac. In analogy to VcINDY, sodium ions are most likely involved in Neu5Ac binding. It has recently been shown that the presence of sodium ions stabilizes the substrate binding site of VcINDY ^28^. As a reverse conclusion, a lower concentration of sodium ions in the cytoplasm would therefore favor the dissociation of the substrate. The P-domain may further enhance this effect by increasing the local substrate concentration at the periplasmic side of the transporter: The relatively confined space of the HiSiaP-QM complex interface has a volume of ∼10^*^10^*^10 Å = 1000 Å^3^. In such a small volume, one Neu5Ac molecule would correspond to a very high local concentration of ∼1 M.

Overall, the results of our study support the current working hypothesis for TRAP transporters and provide experimental evidence that the IFS and OFS of the QM-domains and the closed and open states of the P-domain are indeed conformationally coupled. We show that the HiSiaP CLOSED #1/2, and HiSiaQM IFS/OFS #1/2 proteins are excellent models for mechanistic studies of TRAP transporters. Finally, since all substrate binding proteins share the same architecture, a similar approach may also be useful to further study the mechanism of other SBP-dependent transporters.

## Methods

### Cloning, expression and purification of proteins

The HiSiaQM variants were expressed and purified as published before using a pBADHisTEV vector with N-terminal 10x His-tag and TEV cleavage site ^14^. Mutations were inserted using the method by Liu & Naismith ^29^. The membrane domains were expressed in *E. coli* MC1061 cells in LB-medium (100 μg/ml ampicillin) for 2 h at 37 °C. After harvesting the cells at 4.000 g for 20 min, the pellets were directly used for purification or stored at −80 °C. For purification, the cell pellets were resuspended in 4 times excess buffer A (50 mM KH_2_PO_4_, pH 7.8, 200 mM NaCl and 20% glycerol) and lysed by sonication (40% amplitude, 5 min, pulses 10 s on – 5 s off) on ice. The lysed cells were centrifuged for 1 h and 4 °C at 300.000 g and the pellets were resuspended with a homogenizer in 1.5% (w/v) dodecyl-β-D-maltoside (DDM) supplemented buffer A and incubated overnight under gentle shaking at 4 °C. After an identical ultracentrifugation step, the supernatant was mixed with buffer A equilibrated Ni-NTA agarose beads and incubated for 2 h at 4 °C under gentle shaking. Then, the suspension was loaded on a benchtop column, the flowthrough was discarded, and the beads were washed with 100 ml buffer B (50 mM KH_2_PO_4_, pH 7.8, 200 mM NaCl and 0.035% DDM) with 22 mM imidazole. 15 ml buffer B with 250 mM imidazole was used to elute the protein from the column. For a final SEC purification step, the protein was concentrated to 500 μl (MWCO 100 kDa) and loaded on an equilibrated Superdex 200 increase 10/300 column with buffer B. The eluted SEC fractions were analysed with SDS-PAGE and HiSiaQM containing fractions were concentrated to around 15 mg/ml, flash-frozen and stored at −80 °C.

The HiSiaP proteins were expressed and purified as published before ^9^. The protein was previously cloned into a pBADHisTEV vector, fusing an N-terminal 6x Histag and a TEV cleavage site to the protein. Mutants were prepared using a QuickChange mutagenesis after Liu & Naismith ^29^. *E. coli* BL21 cells were precultured in LB-medium and washed twice with M9 minimal medium. For protein expression, M9 minimal medium was inoculated with bacteria and preincubated 14–16 h at 37 °C before the expression was induced with 500 mg/l L(+)-arabinose. The cells were harvested after 3 h at 37 °C and the pellet was resuspended in 5 times excess of 50 mM Tris (pH 8), 50 mM NaCl. Afterwards the cells were lysed with sonication on ice (40% amplitude, 5 min, pulses 10 s on − 5 s off). After centrifugation for 20 min at 75.000 g the supernatant was filtered and supplemented with equilibrated Ni-NTA agarose beads. After binding for 1 h at room temperature, the suspension was loaded on a benchtop column, the flowthrough was discarded and the column was washed with 100 ml resuspension buffer. The protein was eluted with 500 mM imidazole, concentrated to around 5 ml (MWCO 10 kDa) and loaded on an equilibrated HiLoad Superdex 75 16/600 column (50 mM Tris pH 8, 50 mM NaCl). The eluted fractions were collected and checked with an SDS-PAGE. Protein-containing fractions were combined, concentrated and stored after flash-freezing at −80 °C.

The generation, characterization and sequences of the used VHH_QM_3 was described before ^14^. The protein was expressed with an N-terminal pelB sequence and a C-terminal His_6_-tag in *E. coli* WK6 cells. 1 L of TB medium was supplemented with ampicillin (100 μg/ml) and inoculated with a 25 ml preculture of LB-medium. After incubation at 37 °C to a cell density of 0.6, expression was induced with 1 mM IPTG. After overnight incubation at 30 °C, the cells were harvested by centrifugation at 4.000 g for 20 min. and resuspended in 15 ml extraction buffer (200 mM Tris pH 8.0, 0.65 mM EDTA and 500 mM sucrose) for 1 h at 4 °C under gentle shaking. The resuspended cells were treated with 70 ml of 0.25 times diluted extraction buffer and periplasmic lysis was performed overnight at 4 °C. On the next day, the solution was centrifuged at 8000 g for 40 min, filtered (0.45 μm) and mixed with Ni-NTA beads, equilibrated in 0.25 times extraction buffer. The mixture was incubated for 1 h at 4 °C under gentle shaking and transferred to a bench-top column at room temperature. The flowthrough was discarded, the beads were washed with 50 ml wash buffer (50 mM Tris (pH 7.5), 150 mM NaCl and 10 mM imidazole) and the protein was eluted with 10 ml elution buffer (50 mM Tris pH 8, 150 mM NaCl and 500 mM imidazole). For SEC, the protein was concentrated to 5 ml (MWCO 3 kDa) and loaded on an equilibrated HiLoad Superdex 75 16/600 (10 mM Tris pH 7.3 and 140 mM NaCl). The purification steps and eluted fractions were checked with an SDS-PAGE and protein-containing fractions were combined, concentrated and stored after flash-freezing at −80 °C.

### Fluorophore labelling of HiSiaP and VHH_QM_3

The cysteine mutants of HiSiaP (K254C) and VHH_QM_3 (S85C) were expressed and purified as described above ^14^. Before labelling, each protein was incubated with 1 mM TCEP (Tris(2-carboxyethyl)phosphine) for 30 min at 4 °C and the reducing agent was again removed with a PD Miditrap G-25 column (Cytiva). The eluted protein was directly treated with a 5 times molar excess of AF555 or AF647 maleimide fluorophore (Jena-Bioscience) and incubated for 3 h at 4 °C. Afterwards, the protein was concentrated and washed with a concentrator (MWCO 3 kDa). To remove the remaining unbound label and to check successful labelling, a SEC was performed. The eluted fractions were combined, concentrated and stored after flash-freezing at −80 °C.

For HiSiaP variants CLOSED #1/2 with additional cysteines for stabilization of the closed state, the labelling procedure was slightly changed to avoid labelling of all three cysteines. First, the protein was incubated with Neu5Ac in 1:10 ratio of concentration for 30 min at room temperature. Then, the protein was incubated with H_2_O_2_ (1 mM per 1 mg/ml protein) for 10 min at room temperature and both compounds, the Neu5Ac and H_2_O_2_ were removed (with PD Miditrap G-25 column (Cytiva)). Afterwards, the protein was labelled with the maleimide fluorophore as described above but without the incubation with a reducing agent.

### ITC experiments

ITC experiments were performed on a MicroCal PEAQ device from Malvern Panalytical. The corresponding software was used for design, measurement and analysis of the ITC experiments. The measuring cell was washed with buffer A and loaded with 120 μM HiSiaP solution in buffer A (50 mM Tris pH 8, 50 mM NaCl). The syringe was loaded with 1.2 mM Neu5Ac, dissolved in the same buffer. For each HiSiaP variant (wildtype, CLOSED #1 and CLOSED #2), the experiments were performed as triplicates (data not shown).

### nanoDSF experiments

The nanoDSF measurements were performed on a Prometheus NT.48 from NanoTemper using the corresponding software for measurement and analysis. For each measurement, 25 μl of a 1 mg/ml HiSiaP protein solution was prepared and 10 μl was soaked into a glass capillary (standard capillary, NanoTemper). The start and end temperature were set to 20 °C and 90 °C, respectively, and the heating rate to 1 °C/min. The HiSiaP variants were measured (1) under apo conditions, (2) after 30 min incubation with Neu5Ac at room temperature, (3) after 30 min incubation with Neu5Ac at room temperature and performing a SEC to remove unbound Neu5Ac (see also method preparative SEC runs), and (4) after 30 min incubation with Neu5Ac at room temperature and subsequent 10 min incubation with 1 mM H_2_O_2_ at room temperature and performing a SEC to remove free Neu5Ac and H_2_O_2_.

### X-ray crystallography of HiSiaP

For protein crystallization, HiSiaP was treated with 10 x molar excess of Neu5Ac for 30 min at room temperature and then for 10 min with H_2_O_2_ (1 mM per 1 mg/ml HiSiaP). The solution was cleaned with a Sephadex G-25 column (Cytiva) and the protein was concentrated to 15 mg/ml for crystallization. Sitting drop crystallization attempts were prepared in a 96 Well 2-Drop MRC crystallization plate using the Crystal Gryphon LCP pipetting robot and 0.2 μl of protein solution and 0.2 μl of crystallization screen. The plates were stored at 20 °C in a Rock Imager 1000 (Formulatrix, US) and imaged automatically. Initial hits were observed in PACT screen condition A12 (0.01 M Zinc chloride, 0.1 M Sodium acetate, pH 5.0, 20% w/v PEG 6000) and several hits in a self-designed optimization screen of this condition with small variations in the conditions. Crystals from one optimized condition were harvested, soaked in mother liquor supplemented with 35% glycerol, flash-frozen and stored in liquid nitrogen. Diffraction data of the crystal was recorded at DESY (Deutsches Elektronen Synchrotron, Hamburg) at beamline P13 (λ = 0.826554 Å) using an EIGER 16M detector ^30^. The diffraction data were integrated with autoproc ^31^ and the structure was solved with molecular replacement in PHASER, using the closed-state HiSiaP structure (PDB-ID: 3B50) as model ^32^. The structures were refined and modified with PHENIX ^33^ and COOT ^34^ and validated with MolProbity ^21^.

### Solid supported bilayer preparation

As described previously for TRAP transporter reconstitution into SSB, very small unilamellar vesicles (VSUVs) were prepared from a detergent solution according to Grein et al. and Roder et al. ^14,35,36^. For the bilayer, a lipid mixture of 31.8 mM DOPC and 0.01 mol% TopFluor-PC (both Avanti Polar Lipids) was prepared in chloroform and dried under a nitrogen stream. The lipids were solubilized in 200 μl HEPES buffer (20 mM HEPES (pH 7.4), 150 mM NaCl) with 40 mM Triton X-100. An aliquot of 20 μl was diluted in 200 μl HEPES buffer. 1 μl of HiSiaQM variants at a concentration of 9.2 ng/ml were diluted in 500 μl HEPES buffer and the cysteine variants of HiSiaQM were additionally supplemented with 1 μl of 8 mM H_2_O_2_ and incubated for 10 min at room temperature. Afterwards, 1 μl of this protein solution was added to the lipid mixture. Finally, 200 μl of a 4 mM heptakis(2,6-di-O-methyl)-β-cyclodextrin in water was added and the solution was mixed by vortexing for 2 min. The prepared vesicles were used within 1 h after preparation. 18 × 18 mm coverslips were cleaned overnight in Piranha solution (one-part H_2_O_2_ 30% and two parts concentrated H_2_SO_4_) and rinsed with water. After drying in a nitrogen stream the coverslips were placed into a custom-built chamber with an O-ring as a seal and metal clips to fix the metal insert on top of the coverslip. The bilayers were prepared by adding 400 μl of the VSUVs suspension by filling the well of the sample chamber. Due to electrostatic interactions, a homogeneous bilayer is generally formed after 5 min. Residual vesicles were removed by adding 2 ml of HEPES buffer and removing of only 1 ml of buffer. The chamber was washed 12 times by adding 1 ml HEPES buffer and removing 1 ml. After the last washing step, the final volume in the chamber was 1.4 ml.

### Bilayer binding assay and single molecule imaging

For the VHH_QM_3 staining of HiSiaQM, 2 μL VHH_QM_3-AF555 were added to the surface and incubated for at least 30 min and washed again 5 times as described above by adding 1 ml HEPES buffer and removing 1 ml.

The P-domain variants were diluted 1:20 in HEPES buffer and if necessary for the experiment, the wildtype protein was treated with 10 mM Neu5Ac, incubated for at least 20 min and centrifuged at 14.000 g for 10 min. Wildtype samples without addition of Neu5Ac and samples which were already incubated with Neu5Ac during labelling were treated the same way. To each bilayer 1 μL of this solution was added. The buffer solution in the sample chamber was mixed carefully by pipetting up and down and incubated for 5 min before measurements were started.

Images were acquired at a custom-built, single molecule sensitive, inverted microscope capable of total internal reflection fluorescence (TIRF) microscopy, which was equipped with an sCMOS camera (Prime BSI, Teledyne Photometrics, Tucson, AZ, USA) ^35,37^. Illumination with total internal reflection reduced fluorescence excitation to a thin region at the coverslip surface with the benefit of background suppression from fluorescence outside the illuminated region. The illumination beam angle was adjusted by tilting a collimated laser beam in the object focal plane of the imaging lens, until total reflection at the coverslip/medium interface was reached. This was accomplished by moving the laser beam focus laterally in the back focal plane of the objective, respectively in a conjugated plane located outside the microscope. Using a 63× objective lens with a NA 1.45 (Zeiss) resulted in a pixel size of 103 nm. The focus was always carefully adjusted to the TopFluor-PC signal in the bilayer, which was excited by a laser emitting 488 nm (Cobolt 06-MLD, Hübner Photonics GmbH, Kassel, Germany). The focus was stabilized during the measurements by the definite focus system (Zeiss).

For data acquisition, firstly 1.000 frames using 640 nm or 561 nm (Cobolt 06, Hübner Photonics GmbH) laser excitation for visualizing the P-Domains labelled by AF647 or the Nb3-AF555, respectively, and then 100 frames using 488 nm were acquired. The exposure time was set to 10 ms and only the central 200 × 200 pixels of the camera chip were read out. For each sample 30 measurements were performed and each experiment was repeated for three independent samples.

Processing of the image sequences was performed in Fiji (version 1.52p) ^38^. The 1.000 frames of the P-domain were extracted. Next, the histogram was adjusted to minimum = 0 and maximum = 100 and the background was subtracted using a rolling ball radius of 10. Tracking of single P-domains was performed using the Trackmate plug-in for ImageJ ^39^. For spot detection, the LoG-based detector was chosen. The parameter ‘estimated blob diameter’ was set to 0.75 μm and ‘sub-pixel localisation’ was activated. A threshold of 5 was used for spot filtering. For tracking the ‘Simple LAP Tracker’ was chosen. Gap closing was allowed with a maximum closing distance of 1 μm and a maximum frame gap of two frames. Maximum linking distance was set to 1 μm. Each track was considered as a single interaction of the P-domain with the bilayer.

## Supplementary information

**Supplementary Table 1:** Data collection and refinement statistics.

|  |  |
| --- | --- |
| PDB-ID | 8CP7 |
| Space group | I222 |
| Cell dimensions (Å) |  |
| a, b, c | 84.08, 89.45, 90.5 |
| α, β, γ | 90, 90, 90, |
| Resolution (Å) | 42.04 - 1.9 (1.97-1.9) |
| Wilson B-factor (Å<sup>2</sup>) | 33.96 |
| R<sub>merge</sub> | 0.02891 (0.6132) |
| CC<sub>1/2</sub> | 1.0 (0.721) |
| I/σ(I) | 14.84 (1.14) |
| Completeness | 99.02 (99.43) |
| Redundancy | 2.0(2.0) |
| Unique reflections | 26990(2437) |
| R<sub>work</sub> | 0.2105 (0.3441) |
| R<sub>free</sub> | 0.2589 (0.3640) |
| Average B factor (Å<sup>2</sup>) | 39.37 |
| Protein | 39.47 |
| Ligands | 31.99 |
| Solvent | 38.66 |
| Non-hydrogen atoms | 2635 |
| Protein | 2478 |
| Ligands | 23 |
| solvent | 134 |
| RMS (bonds) (Å) | 0.010 |
| RMS (angles) (°) | 1.01 |
| Ramachandran (%) |  |
| favoured | 99.35 |
| allowed | 0.65 |
| outliers | 0.00 |
| Clashscore | 4.2 |

Values in parentheses refer to the high-resolution shell.

**Supplementary Fig. 1:**
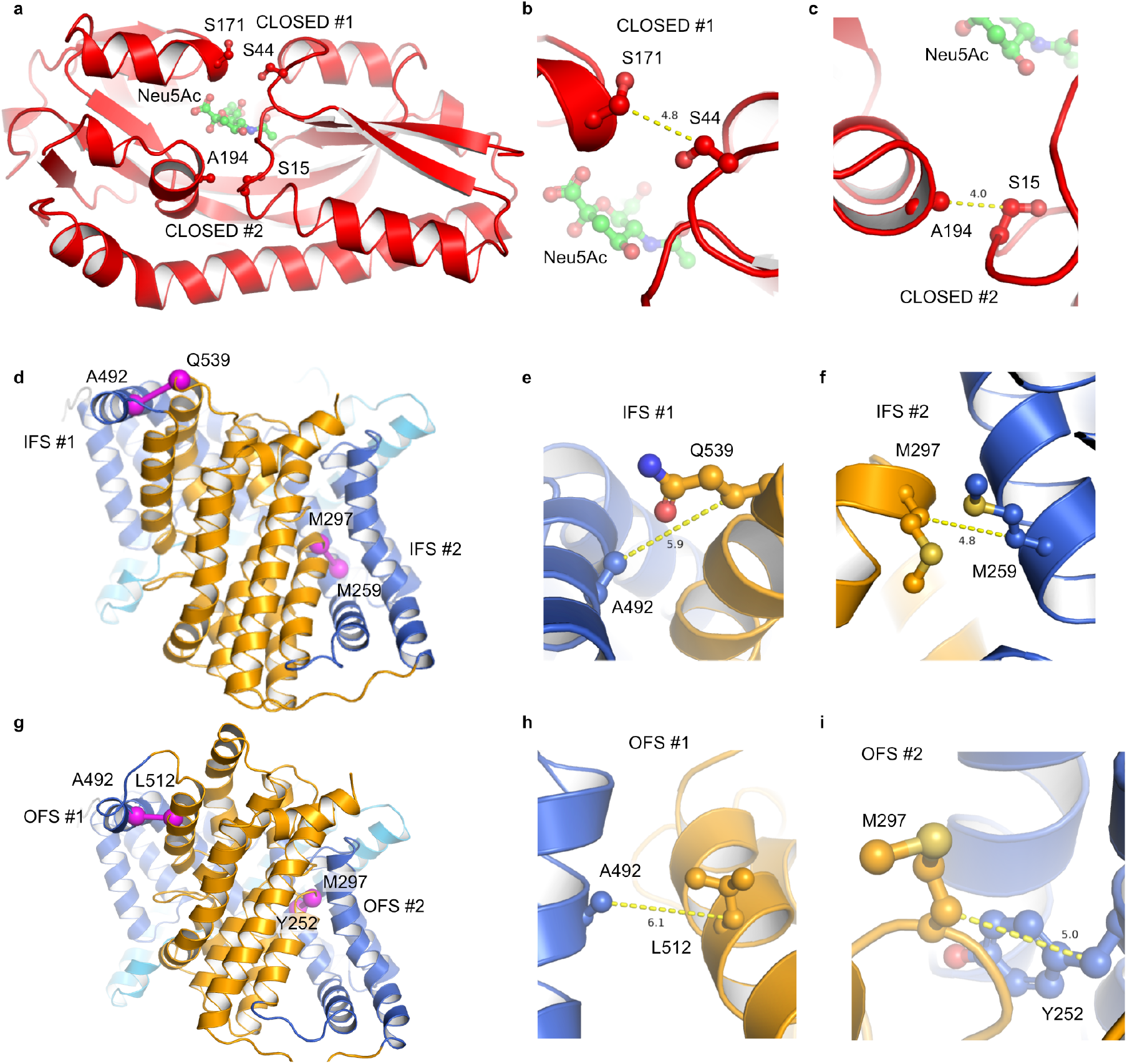
Design of disulfide linked constructs for HiSiaP and HiSiaQM. **a)** Cartoon representation of HiSiaP with Neu5Ac as green sticks and the positions that were mutated to cysteines in CLOSED #1/2 shown as sticks. **b**) Close-up of the CLOSED #1 site. **c**) Close-up of the CLOSED #2 site. **d**) Structure of HiSiaQM in the IFS shown as cartoon model with the positions that were mutated to cysteines shown as magenta spheres. **e**,**f)** Close-up of the IFS#1/2 sites. **g**) Model of HiSiaQM in the OFS shown as cartoon model with the positions that were mutated to cysteines shown as magenta spheres. **h**,**i)** Close-up of the OFS#1/2 sites.

**Supplementary Fig. 2:**
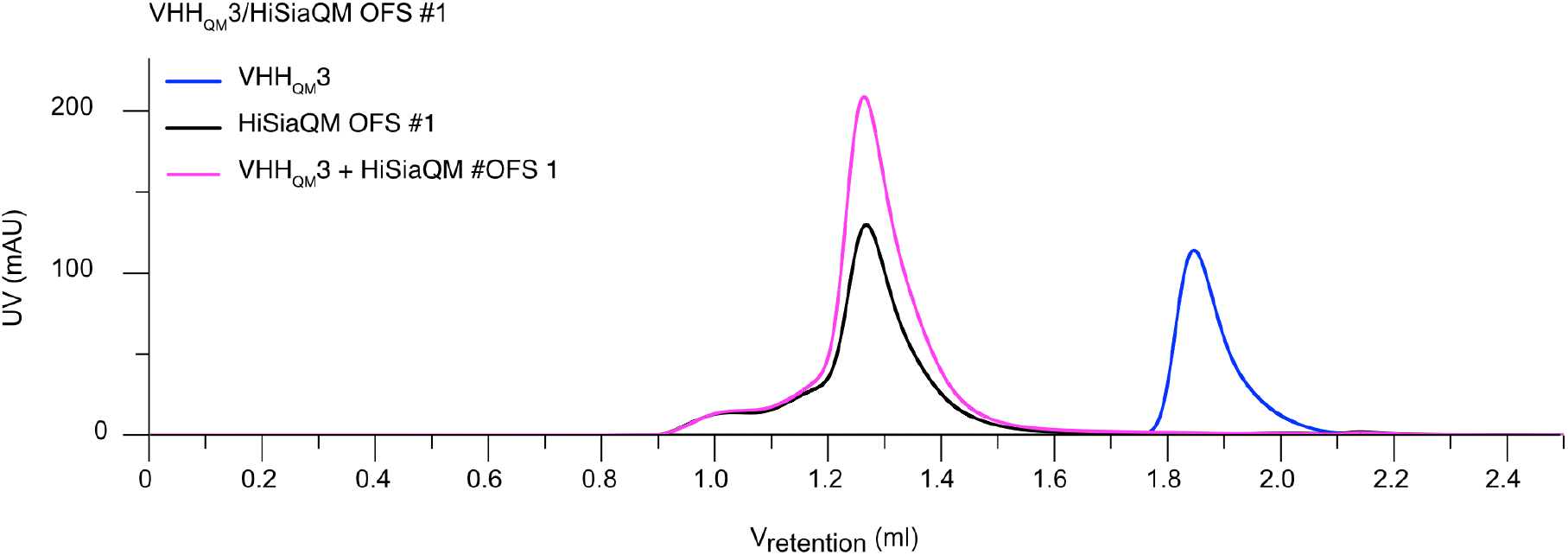
HiSiaQM OFS#1 in detergent binds VHH_QM_3. Three gel filtration experiments with equal amounts of either VHH_QM_3 (blue), HiSiaQM OFS#1 (black) or a mix of both components (magenta). The experiment shows that in DDM detergent, VHH_QM_3 (blue curve) binds to HiSiaQM OFS#1.

**Supplementary Fig. 3:**
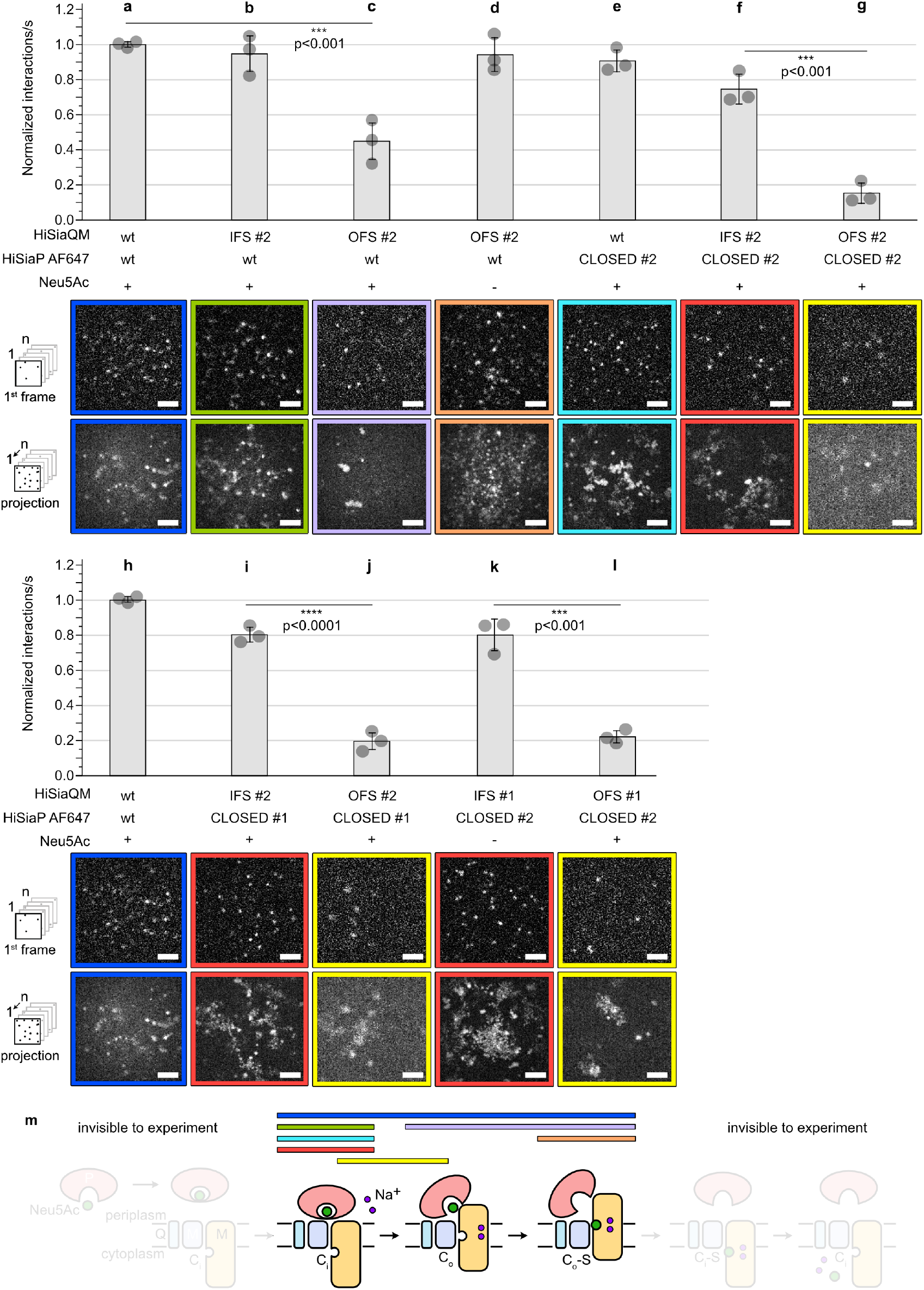
Single molecule TIRF microscopy of trapped TRAP transporter domains. **a-l)** Top: Normalized interactions per second between AF647 labelled HiSiaP variants and the indicated HiSiaQM constructs. Bottom: first frame of an image sequence of a typical set of data and the corresponding maximum intensity projection of the respective image sequence. A movie with all conditions can be found in the supplementary information (Supplementary Movie 1). **m)** In the schematic, colored horizontal bars illustrate the likely state of the transporter observed in the experiments a-g). The statistical significance of differences between selected experiments was assessed by applying a two-sided unpaired Student’s t-test with a 95% confidence interval. The scale bars equal 3 μm.

## Data Availability

The coordinate and diffraction data generated in this study have been deposited in the PDB under accession code 8CP7 [http://doi.org/]. The movie data generated in this study are provided in the Supplementary Information.

## Code Availability

No custom code was used in this study

## Acknowledgements

GH acknowledges funding by the German Research Foundation (DFG), grant number HA 6805/5-1. The synchrotron data was collected at beamline operated by EMBL Hamburg at the PETRA III storage ring (DESY, Hamburg, Germany).

## Author Contributions Statement

GH and MFP conceived this study. MFP, JAR, YK, JPS, GHT, UK & GH planned experiments. MFP, JAR, YK and PH performed experiments. MFP, JAR, YK, PH, JPS, GHT, UK & GH analyzed data. GH and MFP wrote the paper with input from all authors. GH supervised the study and acquired funding.

